# Virtual navigation tested on a mobile app is predictive of real-world wayfinding navigation performance

**DOI:** 10.1101/305433

**Authors:** A. Coutrot, S. Schmidt, L. Coutrot, J. Pittman, L. Hong, J. M. Wiener, C. Hölscher, R.C. Dalton, M. Hornberger, H. J. Spiers

## Abstract

Virtual reality environments presented on tablets and smartphones have potential to aid the early diagnosis of conditions such as Alzheimer’s dementia by quantifying impairments in navigation performance. However, it is unclear whether performance on mobile devices can predict navigation errors in the real world. We compared the performance of 60 participants (30 females, 18-35 years old) at wayfinding and path integration tasks designed in our mobile app ‘Sea Hero Quest’ with their performance at similar tasks in a real-world environment. We first performed this experiment in the streets of London (UK) and replicated it in Paris (France). In both cities, we found a significant correlation between virtual and real-world wayfinding performance and a male advantage in both environments, although smaller in the real world (Cohen’s d in the game = 0.89, in the real world = 0.59). Results in London and Paris were highly similar, and controlling for familiarity with video games did not change the results. The strength of the correlation between real world and virtual environment increased with the difficulty of the virtual wayfinding task, indicating that Sea Hero Quest does not merely capture video gaming skills. The fact that the Sea Hero Quest wayfinding task has real-world ecological validity constitutes a step toward controllable, sensitive, safe, low-cost, and easy to administer digital cognitive assessment of navigation ability.

## 1 Introduction

Virtual reality (VR) provides a powerful mean to study and quantify how humans navigate, because the properties of virtual environment can be completely controlled and repeated across participants. Since the late nineties, it has been a critical tool to understanding how brain regions support navigation and unveiling the structural and functional neural correlates of spatial navigation [1–4]. VR tests of spatial cognition have proved more sensitive in identifying spatial navigation deficits in patient populations compared to more classic visuospatial ‘pencil-and-paper’ tests like the Mental Rotation Test [5]. VR has the added advantage to be a less costly and safer alternative to real-world navigation tests, which are time and space consuming, as well as difficult to administer to a population sometime less able to walk [6]. Until recently, most VR used in research was presented on a desktop display and movement controlled via a joystick or keyboard. Such an interface presents difficulties for older people, less exposed to technology than younger participants [7]. However, with the advent of tablet and smart-phone touch screen mobile devices, older participants have found engaging in VR tasks much easier and intuitive than with desktop computers [8, 9]. As a consequence, mobile devices has recently been used in several fields such as neuropsychological assessment [10], stroke rehabilitation [11] and mental health [12]. We recently developed a VR navigation task for mobile and tablet devices — Sea Hero Quest — with the aim that this may provide an early diagnostic tool for Alzheimer’s Disease (AD) [13]. For this test to be useful it is important that it has real-world validity, with errors on the VR task predicting errors in real-world navigation experience.

Past research comparing navigation in real and VR environments has generally found a good concordance in performance across both environments in the normal population [14–19], in younger and older age groups [20], in individuals with brain injury [21, 22], in chronic stroke patients [23], and in patients with Mild Cognitive Impairment (MCI) or early AD [20, 24], for reviews see [25, 26]. However, this consistency seems to be modulated by the type of spatial navigation task, as a previous study showed that performance in real life and virtual environments were similar for tasks such as landmark recognition or route distance estimate, but different for pointing to the beginning and endpoint of the route, or drawing a map of the route [27].

Most prior studies comparing VR and real-world navigation performance have used desktop VR or immersive VR to simulate environments, and paper and pencil tests such as line orientation, road map, or delayed recall when assessing ‘real-world navigation behavior’. A few studies made use of actual navigation tasks but often in a limited spatial range, like the lobby of a hospital [20]. A notable exception is [32], where the authors tested 978 military college students on a 6 km orienteering task and replicated many laboratory-based findings, including gender differences. However the authors did not test their participants in a VR task and were thus unable directly compare the two environments.

Numerous studies found a male advantage for navigation in VR tasks [13, 28–31], but only a few looked for gender differences in real-world navigational tasks [32, 33]. This led some authors to suggest that previously reported gender differences in spatial ability may be driven by familiarity with technology, men being more comfortable with virtual tasks than women who are sometimes less exposed video games [34].

Here, for the first time we directly compared in a within-subject design the spatial navigation performance measured on a mobile device with our Sea Hero Quest virtual tasks, and in a large-scale real-world environment covering a whole neighborhood of London (Covent Garden, South of the British Museum) and of Paris (South of the Montparnasse cemetery). We designed the real-world counterparts of the Sea Hero Quest wayfinding and path integration tasks, which are known to tap into different cognitive processes [13]. We hypothesized that performance in the real world in both cities will significantly correlate with performance in the virtual environment. Based on our original mobile-based results [13] and on previous navigation studies highlighting gender differences in the real world [32], we hypothesized that males will perform better than females in both environments. Finally, we predicted that familiarity with video games will not influence performance in either environment because navigation skill requires different abilities to manipulating the controls in a video game. In particular, the correlation between the real-world wayfinding performance and the performance at the first training Sea Hero Quest level – where no spatial ability is required – should be null. Comparing this study to the original large dataset would enable testing whether our results hold true not simply in small cohorts but on a population level.

## 2 Methods

Participants were tested on specific levels from Sea Hero Quest [13] on a tablet, and then on equivalent tasks in the real world, see Fig 1. Participants were also asked to answer a few demographic questions. We first ran this experiment in London in summer/fall 2017. We then replicated it with a different team in Paris in spring 2018. The whole experiment lasted around three hours.

**Figure 1.**
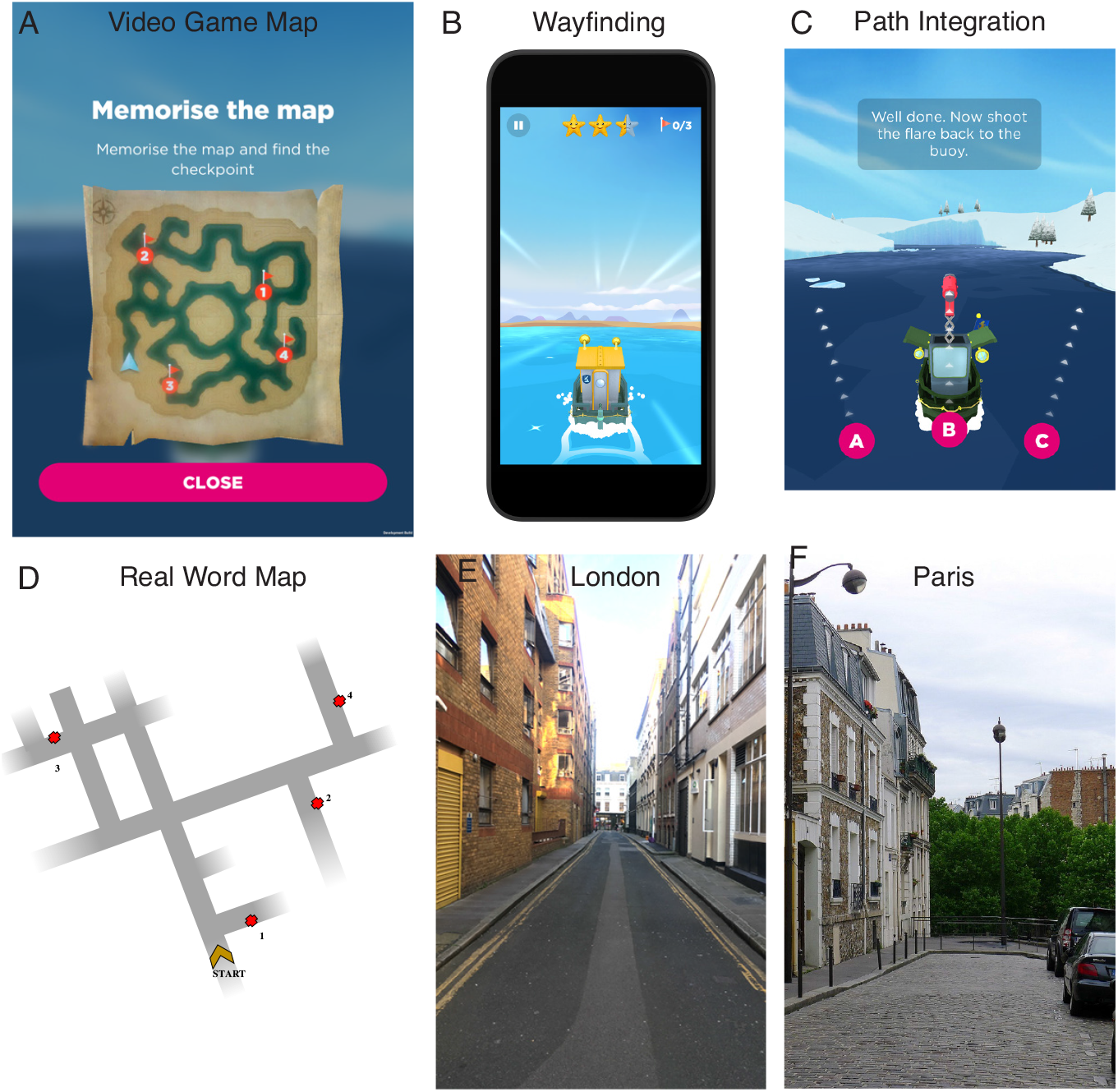
Task in real world (bottom row) vs virtual environment (top row). (A-B) Wayfinding task in the video game: participants had to memorize a map and navigate as fast as possible toward an ordered set of goals. Participants played Sea Hero Quest on a tablet. (C) Path Integration task in the video game: participants had to navigate in a maze until they find a flare and shoot it back toward their starting position. (D) Wayfinding task in the real world. Identical as the virtual task, but takes place in the streets of (E) London and (F) Paris. All other maps are displayed in supporting figures S1 to S4.

### 2.1 Participants

#### In London

We tested 30 participants (15 males), aged 18-30 y.o. (M= 21.96, s.d.= 2.55). Data from one participant was missing due to a technical problem. Real-world wayfinding data from six participants (three males and three females) were discarded due to GPS recording issues. Participants had normal or corrected to normal vision and gave their written consent to participate. Path integration data was not collected for the first 11 participants as this task was not yet implemented. Participants received 3 class credits or £20 for their participation.

#### In Paris

We tested 30 participants (15 males), aged 18-35 y.o. (M= 24.27, s.d.= 3.31). Real-world wayfinding data from four participants (one male and three females) were discarded due to GPS recording issues. Participants had normal or corrected to normal vision and gave their written consent to participate. Participants received 30 euros for their participation.

### 2.2 Virtual tasks

We devised a mobile video game designed to measure human spatial navigation ability through gameplay - Sea Hero Quest (SHQ, www.seaheroquest.com). This video game involves navigating a boat in a virtual environment (lake or river networks) and has been extensively described in [13]. It features two main tasks, which has been designed to tackle different aspects of spatial navigation.

#### 1-Wayfinding

Participants were required to view a map displaying current position and goal locations to find (Fig 1A-B). Participants could study the map without time restrictions and had to navigate to the goal locations in the order indicated, e.g. goal 1 must be found first, then goal 2, etc. Goals were buoys with flags marking the goal number. The task is complete when all goals have been located. If the participant takes more than a set time, an arrow indicates the direction along the Euclidean line to the goal to aid navigation. On basis of the data from the online version, we selected a subset of 5 of the total 75 levels in the game that varied in difficulty: instruction level 1, and wayfinding task levels 6, 11, 16, and 43 (see S1 Fig). The helping arrow appeared after 80 s in level 1, 70 s in level 6, 80 s in level 11, 80 s in level 16 and 200 s in level 43. Performance was quantified with the Euclidean distance travelled in each level (in pixels). The coordinates of participants’ trajectories were sampled at Fs = 2 Hz. We summed the distance travelled over levels 6 to 43. We did not include level 1 because it did not require any spatial ability (the goal was visible from the starting point) and was only designed to assess participants’ ability to learn to control the boat.

#### 2-Path Integration

Participants were required to navigate along a river with bends until they find a flare and shoot it back toward their starting position. Participants could choose among three directions, as shown in Fig 1C. We selected a subset of 5 levels that varied in difficulty: level 14 had one bend, level 34 three, level 44 two, level 54 four and level 74 five, see S4A Fig. Performance was measured with the number of stars obtained by the player. Stars were awarded based on participant’s choice between 3 proposed directions: 3 stars for the correct answer (their starting point), 2 stars for the second closest direction, and 1 star for the third closest direction.

### 2.3 Real-world task

#### 1-Wayfinding

The real-world wayfinding task consisted of 6 wayfinding trials which varied in difficulty in terms of the number of streets to be navigated, the number of goals and the relative location of the goals to each other. Each trial was located in a different street network in South of the British Museum in London (Covent Garden area) and South of the Montparnasse cemetery in Paris. We chose less busy streets to avoid traffic and made sure the participants were not familiar with them. Before each trial, participants were shown a map that only indicated the facing direction, the network of the local streets and the location and the order of the goals (in London see S2 Fig, in Paris see S3 Fig). Maps were displayed on a tablet (Ipad MP24B/A, 9.7 inches). The goals were doors and gates with distinct features (e.g. specific colour, size, or material). Participants had up to 1 min to memorize the map (maximum length for Sea Hero Quest) before walking to locate the goals. During navigation they were provided with colour photographs of the goal. Based on pilot testing we set specific time limits for each route. We chose these time limits to allow for a few mistakes at a reasonable walking pace. Pilot testing indicated that if participants required any longer than that these times they were likely guessing and had failed to remember the goal locations or street layout. If participants reached the limits of the defined region set by the experiment they were told by the experiment they had reached the edge of the search area and should turn back. In London, route one: 6 minutes, route two: 6 minutes, route three: 6:30 minutes, route four: 6:30 minutes, route five: 12 minutes, route six: 14 minutes. In Paris, route one: 5 minutes, route two: 8 minutes, route three: 8 minutes, route four: 9 minutes, route five: 16 minutes, route six: 20 minutes.

The coordinates of participants’ trajectories were sampled at Fs = 1 Hz with the experimenter’s smartphone GPS via the Beeline app. We visually inspected all recorded GPS trajectories to deal with potential losses of signal. For losses of signal where the participant did not make any turn, we linearly interpolated between the first and the last missing points. When we couldn’t reconstruct the trajectory because the participant changed direction during the loss of signal, we discarded the data (3.3% of the trials in Paris, 2.8% in London). Performance was quantified with the Euclidean distance travelled in each route (in meters). To take into account the fact that some participant did not finish some routes, we divided this distance by the number of goals reached by the participant plus 1. We added 1 to cope with cases where the participant didn’t reach any goal (this only happened once). We refer to this as the metric normalized distance, and summed it over routes 1 to 6.

#### 2-Path Integration

The real-world path integration task consisted of 4 path integration trials which varied in difficulty in terms of the number turns they featured (1, 2, 3 and 4 turns, see S4B Fig). To avoid familiarity effect, path integration routes were chosen not to intersect with any wayfinding route. Participants were required to follow the experimenter to an endpoint where they were asked to point back toward the starting point. We used a numeric compass to precisely record the direction. Performance was defined as the inverse of the angle between the direction pointed toward by the participant and the ground-truth, in degrees. We then summed the absolute values of the path integration error angles.

## 3 Results

Table 1 reports the difficulty of each wayfinding route in the real-world task as the percentage of goals reached by the participants before the time limit. This goes from 99% in London (100% in Paris) for route 1 down to 74% in London (84% in Paris) for route 6. Table 1 also reports the difficulty of each wayfinding level in the video game task as the percentage of participants that managed to complete the level before the helping arrow appeared. 100% participants successfully completed the first training level while only 47% participants in London (40% in Paris) completed level 43. Interestingly, level 11 seemed harder than level 16, which might be due to level 11 requiring participants to turn back to meet the goals in order, which was not the case in level 16 (see S1 Fig).

**Table 1.**
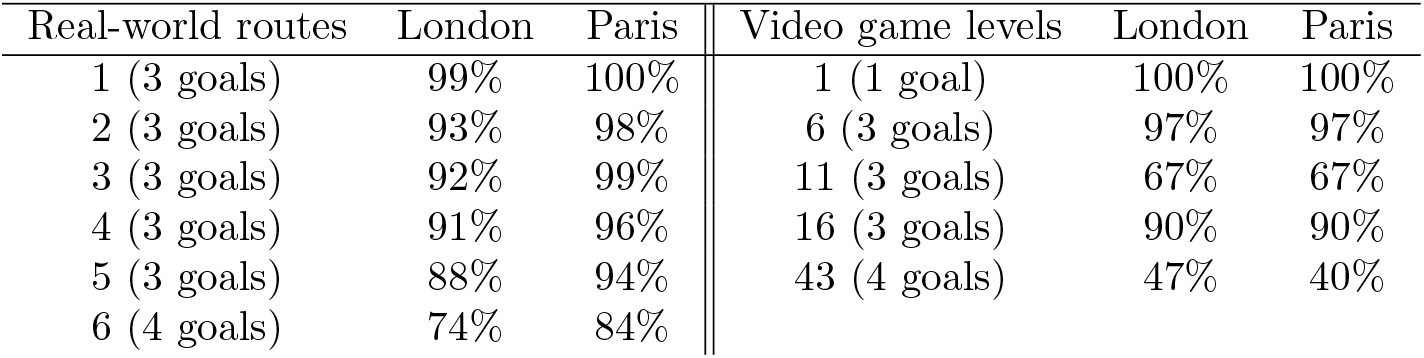
Difficulty of the virtual and real-world wayfinding task. Left: Real-world wayfinding task. For each route, average percentage of goals reached by the participants in London and Paris within the time limit. Right: Video game wayfinding task. For each level, average percentage of participants that completed the level before the helping arrow appeared.

| Real-world routes | London | Paris | Video game levels | London | Paris |
| --- | --- | --- | --- | --- | --- |
| 1 (3 goals) | 99% | 100% | 1 (1 goal) | 100% | 100% |
| 2 (3 goals) | 93% | 98% | 6 (3 goals) | 97% | 97% |
| 3 (3 goals) | 92% | 99% | 11 (3 goals) | 67% | 67% |
| 4 (3 goals) | 91% | 96% | 16 (3 goals) | 90% | 90% |
| 5 (3 goals) | 88% | 94% | 43 (4 goals) | 47% | 40% |
| 6 (4 goals) | 74% | 84% |  |  |  |

Table 2 reports the difficulty of each path integration route in the real-world task in Paris as the mean absolute error angle, and the difficulty of each path integration level in the video game task in Paris as the percentage of correct answers (i.e. the percentage of participants that received three stars). London data is not displayed since the first 11 participants were not tested on path integration, as this task was not designed yet.

**Table 2.** Difficulty of the virtual and real-world path integration task in Paris. Left: Real-world path integration task. For each route, average error angle (*M* ± *SE*, in degree). Right: Video game path integration task. For each level, average percentage of participants that received three stars (correct answers).

| Real-world routes | mean error angle | Video game levels | % of correct answers |
| --- | --- | --- | --- |
| 1 (1 turn) | $19.8^\circ \pm 4.5$ | 14 (1 turn) | 87% |
| 2 (2 turns) | $21.7^\circ \pm 2.8$ | 34 (2 turns) | 43% |
| 3 (3 turns) | $19.5^\circ \pm 2.4$ | 54 (3 turns) | 70% |
| 4 (4 turns) | $25.6^\circ \pm 4.4$ | 44 (4 turns) | 73% |
|  |  | 74 (5 turns) | 20% |

### Correlation between performance in real world and virtual environments

To visualize participants’ raw data in real world and virtual environments we created a video showing on the left side the trajectories of participants in London’s route 6 and on the right side the trajectories of the same participants in the 43rd level of Sea Hero Quest (S1 Vid). The relationship between the wayfinding performance in the real world and in the video game is shown Fig 2A (in London) and 2B (in Paris). One can notice a few outliers in the upper right corner of the scatter plots. Traditional Pearson’s correlation measure is known to be highly sensitive to outliers, which can severely bias the estimation of the strength of the association among the bulk of the points [35]. To deal with this, we used skipped-correlation, which protects against outliers by taking into account the overall structure of the data [36]. Outliers are detected using a projection method, removed, and Pearson’s correlation is computed using the remaining data. Hence, Pearson’s skipped correlation is a direct reflection of Pearson’s r. We used an implementation of this algorithm available in a free toolbox [37]. The detected outliers are tagged in Fig 2A-B with black edges, and discarded from further analysis. 95% Confidence Intervals (CI) were computed via bootstrap: pairs of observations were resampled with replacement and their correlation values obtained and sorted. Values between the 2.5 and 97.5 percentiles yielded the 95% CI. Skipped correlation were significant both in London (*r* = 0.46, 95% CI = [0.14, 0.68], *p* = 0.01) and in Paris (*r* = 0.57, 95% CI = [0.37, 0.76], *p* = 0.001).

**Figure 2.**
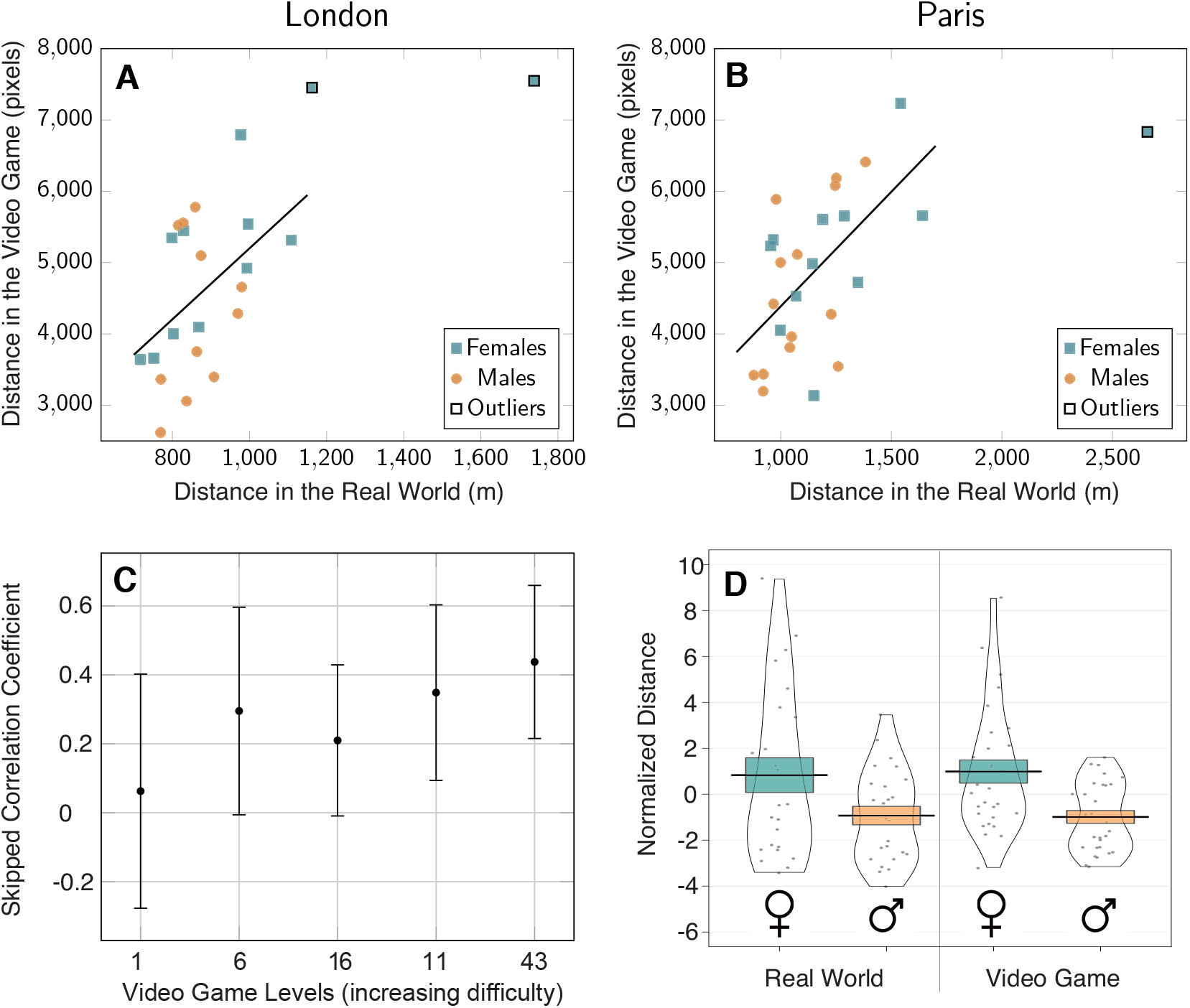
Spatial ability at a wayfinding task in real world vs virtual environment. (A) Correlation between the distance navigated in the video game and real-world wayfinding task in London (skipped Pearson’s *r* = 0.46, *p* = 0.01) and (B) in Paris (skipped Pearson’s *r* = 0.57, *p* = 0.001). Outliers have been determined with skipped-correlation. (C) Skipped correlation coefficients between the distance navigated in each video game level and the total distance navigated in the real world. Video game levels are sorted by increasing difficulty according to Table 1. (D) Gender differences at the wayfinding task in the video game (right) and in the real world (left). Black dots represent individual data points. Error bars represent standard errors.

To confirm that our virtual task captured participants’ wayfinding ability, we checked whether the strength of the correlation between performance in real world and in the video game was modulated by the difficulty of the virtual task. Under this hypothesis, the correlation should be null between real world and level 1 performance, since level 1 is a training level where no spatial ability is required: the end goal is visible from the starting point. The correlation should then increase with the difficulty of the level. We broke down the global correlation score for each Sea Hero Quest level, comparing participants’ performance in real life with the distance they travelled in each level. To increase the sample size, we combined the data recorded in London and in Paris. In order to take into account the difference in route length between cities (the task in Paris being slightly longer than the one in London), we calculated zscores of the performance for each level before combining the data of the two cities. In the following we work with this cross-city normalized metric, called Normalized Distance. The skipped correlation coefficient between the Normalized Distance in the real world and in level 1 is close to 0, confirming that this instruction level does not measure spatial ability. We sorted the game levels by increasing difficulty based on the results of Table 1: level 1, 6, 16, 11 and 43. As shown in Fig 2C, the skipped correlation coefficient increases with level difficulty, from r = 0.06 in level 1 to r = 0.44 in level 43.

The skipped correlation between real world and video game path integration score in Paris was not significant, r = −0.23, 95% CI = [-0.51, 0.08], p = 0.11. However, the sign of the correlation is logical since higher scores mean better performance in the game (number of stars) but not in the real-world task (error angle). In London, the first 11 participants were not tested on path integration as this task was not yet designed. The skipped correlation based on the other 19 participants is consistent with Paris data: r = −0.40, 95% CI = [-0.73, 0.07], p = 0.05. However, correlational analyses with small sample size can lead to strongly biased correlation estimates and this result should be considered with caution.

### Gender differences

For the wayfinding task, we found that in both environments male participants had an advantage, although smaller in the real world (Fig 2D). In the real world, Cohen’s *d* = 0.59, 95% CI = [0.02 1.15], t(47) = −2.08, p = 0.04; in the video game, Cohen’s *d* = 0.89, 95% CI = [0.35 1.42], t(56) = −3.43, p = 0.001.

For the path integration task, we did not find a significant gender difference. However, the tendency was similar to the wayfinding results, with male having a small advantage in both environments, smaller in the real world. In the real world, Cohen’s *d* = −0.10, 95% CI = [-0.79 0. 60], negative values correspond to a male advantage; in the video game, Cohen’s *d* = 0.29, 95% CI = [-0.40 0.99], positive values correspond to a male advantage. The gender effect size is much smaller for path integration than for wayfinding and is not significative, as shown by the wide 95% CIs.

### Influence of familiarity with video games

On average, females played video games 2.99 ± 8.38 hours per week and males played 2.95 ± 4.21 hours per week. To control for the influence of familiarity with video games on real-world and virtual spatial abilities, we computed a multiple linear regression to predict performance based on gender and on the time participants spent playing video game (VGT), in hours per week.

With normalized distances recorded in the real-world wayfinding task, gender was a significant predictor (*t*(47) = —2.22, *p* = 0.03), but not VGT (*t*(47) = 0.31, *p* = 0.76). Similarly, with normalized distances recorded in the virtual wayfinding task, gender was a significant predictor (*t*(55) = —3.44, *p* = 0.001), but not VGT (*t*(55) = — 0.99, *p* = 0.32).

With error angles recorded in the real-world path integration task, gender was not a significant predictor (*t*(55) = —0.65, *p* = 0.52), nor was VGT (*t*(55) = 1.09, *p* = 0.28). Similarly, with normalized flare accuracy recorded in the virtual path integration task, gender was not a significant predictor (*t*(55) = —0.17, *p* = 0.87), nor was VGT (*t*(55) = 0.09, *p* = 0.92).

### Correlation between wayfinding and path integration

The skipped correlation between path integration and wayfinding scores is not significant in the real world (Paris data): r = −0.06, 95% CI = [-0.34 0.44], nor in the virtual environment: r = −0.18, 95% CI = [-0.49 0.15]. Higher scores mean better performance in the path integration task in the game (number of stars) lower score mean better performance in the path integration task in the real-world task (error angle), and in the wayfinding tasks in both environments (distance).

### Comparison to the population-level original dataset

To check whether the 60 participants we recruited for this experiment were representative of the much larger dataset recorded with the mobile version of Sea Hero Quest [13], we plotted in Fig 3 the performance of the participants at this study (vertical red dotted lines) against the corresponding distribution of the performance of the French and British Sea Hero Quest players from the original dataset. Since the number of players per level drops rapidly, we focused on level 11 to maximise the difficulty / number of players ratio (N = 78,724, see [13] for full data). Fig 3 clearly shows that the performance of the participants recruited for this study closely follows the performance distribution of the original dataset. Gender differences followed the same direction in this study (Cohen’s d = 0.81, 95% CI = [0.28 1.33]) as in the subsample of the original dataset (Cohen’s d = 0.40, 95% CI = [0.39 0.42], positive values correspond to a male advantage.

**Figure 3.**
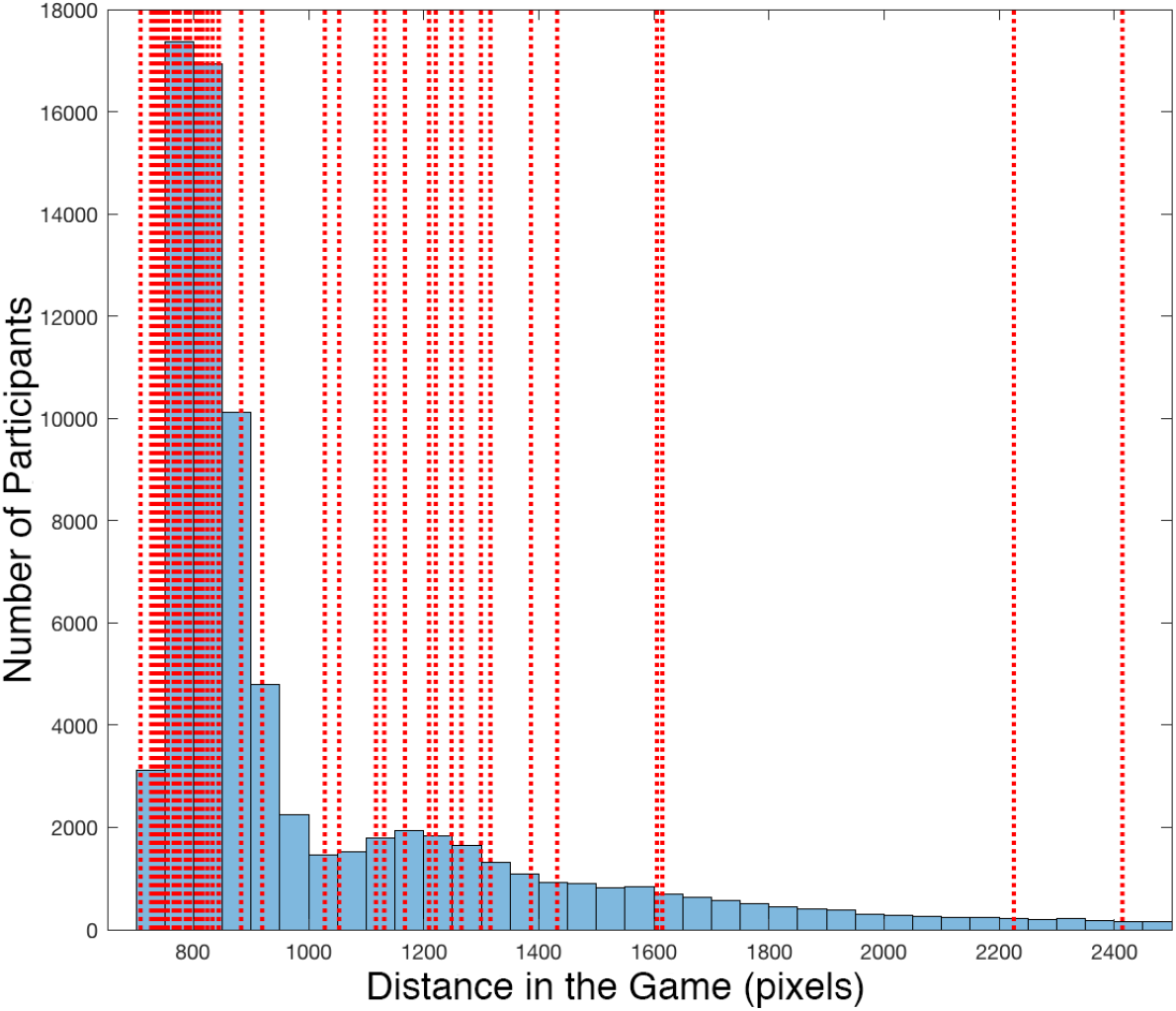
Distance in the Video Game (level 11) of the French and British participants tested in the original Sea Hero Quest database (N = 78,724, blue histogram), see [13] for full data. The red dotted vertical lines represent the performance of the participants recorded in the current study.

## 4 Discussion

We report evidence that wayfinding navigation performance on a mobile app-based VR navigation task (Sea Hero Quest) is significantly correlated with performance in a real-world city street wayfinding task. We directly compared participants performance at a subset of Sea Hero Quest wayfinding levels with their performance at an equivalent task in the Covent Garden area, London. We found a strong correlation between the distance participants travelled in the video game (in pixels) and in the real-world street network (in meters, measured by a GPS device). We replicated this result with another set of participants in the Montparnasse area, Paris. The high similarity of the results in the two cities is a strong indicator of the robustness of the results presented above. Our findings are consistent with a number of studies that showed that spatial navigation assessment in a desktop VR [15, 16, 18, 20, 22, 23, 27] and immersive VR [17, 38] environments transferred well to the real world, and extend them to tablet device presentation and real-world spatial task spanning complex street networks. The similarity both in term of performance and of gender difference of this study with the original large scale Sea Hero Quest study [13] suggests that our findings hold true not simply in small cohorts but on a population level.

We found a significant male advantage in the wayfinding task in both real world and virtual environments, although weaker in the real world. This difference in effect size between the two environments couldn’t be explained by males being more familiar with video games, as suggested in a previous study [34] for three reasons. First, our male and female participants reported playing video games the same average duration per week (2.95 vs 2.99 hours per week). Second, when using gender and time playing video games (VGT) as covariates in a linear model to predict wayfinding performance in the real world (resp. in the virtual environment), gender came out as a significant predictor, but not VGT. Third, we found a very weak correlation coefficient (skipped Pearson’s r = 0.06) between the performance at the real-world wayfinding task and at the first training Sea Hero Quest level, which did not require any spatial ability (the endpoint being visible from the start). The strength of the correlation increased with the difficulty of the video game level (up to r = 0.44 in level 43), indicating that Sea Hero Quest does not merely capture video gaming skills. The discrepancy with [34] might stem from the difference between tasks, Richardson et al.’s task being closer to the path integration task than to the wayfinding task discussed in this paragraph.

We did not find a significant correlation between the performance at the real world and the Sea Hero Quest path integration task. At least three reasons could account for this null result. First, as mentioned in the introduction, the consistency between spatial navigation ability in the real world and in a virtual environment task depends on the type of navigational task [27]. In particular, the aforementioned study reported a poor concordance for a task involving pointing to the beginning and endpoint of the route, which is quite close to our path integration task. This hypothesis is consistent with the weak correlations we found between wayfinding and path integration performances, both in the real world and in the virtual environment: the two tasks involve different cognitive processes, which don’t generalize similarly from one environment to the other. Second, this null result could be caused by the low sensitivity of our virtual path integration task. Indeed, while in the real-world performance was a continuous variable defined as the inverse of the error angle, in Sea Hero Quest it could only take three values: one, two, or three stars. This ternary metric might not be sensitive enough to capture subtle differences in the moderate sample size used in this study (60 participants), unlike the original Sea Hero Quest study on mobile and tablet (2.5m participants) [13]. Third, one could argue that path integration in a city is different from path integration in a controlled virtual environment, as there are environmental structures (e.g. street grid) and landmarks (e.g. buildings), which may help to judge distances and directions. This would explain the small difference in mean error angle captured by the real-world path integration routes between supposedly easy (one turn) and difficult (four turns) routes, see Table 2.

Altogether, these results constitute a step toward the ability to remotely test people. This is particularly valuable when certain categories of the population have difficulties in mobility, like older people. Currently our results focused on young university students, and it will be useful to extend to a broader population including elderly participants. Spatial ability assessment provides the potential to act as an early stage diagnostic tool for Alzheimer’s dementia (AD), because spatial disorientation is one of the earliest symptoms [24, 39–43]. Currently there is no standardized test for navigation deficits with AD patients, as diagnostics measures are still focused on episodic memory deficits, despite their low sensitivity and specificity for identifying at-risk individuals [26]. Sea Hero Quest wayfinding task having real-world ecological validity holds future promise for controllable, sensitive, safe, low-cost, and easy to administer digital cognitive assessment.

## Data Availability

All the data supporting this study will be made available on the same server as the main Sea Hero Quest dataset [13]. This server is currently under active development. In the meantime data will be provided upon request to the lead author.

## Author Contributions

Conceptualization: HS, MH Data

Curation: AC, SS, LC, JP, LH

Project Administration: AC, HS

Supervision: AC, HS

Investigation: AC, SS, LC, JP, LH

Methodology: AC, HS, SS, LC

Validation: AC

Visualization: AC

Writing – Original Draft Preparation: AC, HS

Writing – Review & Editing: AC, JW, CH, RD, MH, HS

